# Why Are CD8 T Cell Epitopes of Human Influenza A Virus Conserved?

**DOI:** 10.1101/408880

**Authors:** Zheng-Rong Tiger Li, Veronika I. Zarnitsyna, Anice C. Lowen, Daniel Weissman, Katia Koelle, Jacob E. Kohlmeier, Rustom Antia

**Author notes:** Address correspondence to Rustom Antia or Veronika I. Zarnitsyna.

## Abstract

The high-degree conservation of CD8 T cell epitopes of influenza A virus (IAV) may allow T cell-inducing vaccines effective across different strains and subtypes. This conservation is not fully explained by functional constraint, since additional mutation(s) can compensate the replicative fitness loss of IAV escape-variant. Here, we propose three additional mechanisms that contribute to the conservation of CD8 T cell epitopes of IAV. First, influenza-specific CD8 T cells may protect predominantly against severe pathology rather than infection and may only have a modest effect on transmission. Second, polymorphism of human MHC-I gene restricts the advantage of an escape-variant to only a small fraction of human population, who carry the relevant MHC-I alleles. Finally, infection with CD8 T cell-escape-variants may result in compensatory increase in the responses to other epitopes of IAV. A combination of population genetics and epidemiological models is used to examine how the interplay between these mechanisms affects the rate of invasion of IAV escape-variants. We conclude that the invasion of an escape-variant will be very slow with a timescale of decades or longer, even if the escape-variant does not have a replicative fitness loss. Our results suggest T cell-inducing vaccines may not engender the rapid evolution of IAV and serve as a foundation for future modeling works on the long-term effectiveness and impacts of T cell-inducing influenza vaccines. (Word count: 221)

**Importance:** Universal influenza vaccines against the conserved epitopes of influenza A virus have been proposed to minimize the burden of seasonal outbreaks and prepare for the pandemics. However, it is not clear to which extent the T cell-inducing vaccines will select for viruses that escape the T cell responses. Our mathematical models suggest how the nature of CD8 T cell protection contributes to the conservation of the CD8 T cell epitopes of influenza A virus. Also, it points out the essential biological parameters and questions that need addressing by future experimental works. (Word count: 91)

## 1 Introduction

Seasonal influenza is a major public health concern, causing about 410,000 deaths worldwide annually (1). The inactivated vaccine currently in use requires an antigenic match between the vaccine and circulating strains, so that the vaccine-induced antibodies can block viral entry and exit by binding to the antigenic sites, or epitopes. Most of the antibody epitopes reside on hemagglutinin (HA) and neuraminidase (NA), both of which vary largely across strains and subtypes of influenza A virus (IAV). Antibodies induced by exposure to earlier IAV strains drive the selection of antibody-escaping mutants (2). These new drifted strains of the virus are responsible for seasonal outbreaks of influenza. In addition to gradual changes caused by antigenic drift, larger antigenic shifts may occur when new antigenic subtypes (e.g., H1, H3, H5, H7) emerge from zoonotic reservoirs into the human population. Both antigenic drift and antigenic shift necessitate frequent updating of the influenza vaccine. New approaches that focus on antibodies against the conserved regions of HA or CD8 T cells specific to the conserved epitopes on interior proteins have been proposed for the development of vaccines that have broad efficacy and ideally confer ‘universal’ protection against all IAV subtypes (3, 4, 5). Whether IAV is likely to evolve and escape the vaccine-induced CD8 T cell immunity is an essential question for the development of ‘universal’ CD8 T cell-inducing vaccines. Therefore, in this paper, we investigate the reasons why CD8 T cell epitopes of IAV are so conserved.

CD8 T cells detect and kill virus infected cells by recognizing short viral protein-derived peptides (epitopes) bound to major histocompatibility complex class I proteins (MHC-I) on cell surface. While CD8 T cells are not considered to generate ‘sterile immunity’ that prevents infection, they can reduce the severity of disease and potentially viral transmission (6, 7). In addition, they may provide broad protection since CD8 T cell epitopes of influenza virus are largely conserved among drifted strains within one subtype and across different subtypes (7, 8, 9). The conservation of CD8 T cell epitopes is consistent with the observation that internal viral proteins (such as nucleoprotein (NP) and matrix-1 protein (M1)), which harbor the majority of CD8 T cell epitopes, have a much lower substitution rate than HA and NA, which are the targets of antibody responses (10). Also, within NP and M1, the epitope regions have less sites of *dN/dS* > 1 compared to the non-epitope regions (11). Unlike the highly variable antibody epitopes, to date only 6 out of 64 CD8 T cell epitopes in human IAV have been found to have mutations that allow escape from CD8 T cell responses (9, 12).

Why are CD8 T cell epitopes of human influenza virus conserved? The functional constraint hypothesis proposes that CD8 T cell epitopes are on the protein regions that do not tolerate changes in the amino acid sequence (13, 14). In this hypothesis, non-synonymous mutations in the epitope region totally or partially diminish protein function and result in viruses that incur a cost at the level of replication. In support of this, several studies have revealed that a single amino acid change at some residues within the NP or M1 can reduce the replication of the virus (13). A more recent study has shown that epistatic interactions may stabilize mutations and permit previously inaccessible destabilizing mutations (15). While the epistatic interactions greatly ameliorate the fitness cost, it would slow the rate at which a competitive mutant is generated. In addition, at least two other factors may contribute to the conservation of CD8 T cell epitopes. First, while CD8 T cells protect the hosts from severe pathology, they might not greatly reduce virus transmission from an infected individual to new hosts; namely, the viruses are under no or little selection pressure. Second, a mutant that has one or more mutated CD8 T cell epitopes may have selective advantage only in the hosts whose MHC-I present the wild-type form of CD8 T cell epitope; thus, polymorphism of MHC-I genes limits the selective advantage of a mutant in a fraction of host population. In this paper, we use simple mathematical models to explore how the interplay between above-mentioned factors affects the rate of invasion of a mutant, which has a mutated epitope that is not detected by CD8 T cells specific for the wild-type epitope. For brevity, the two virus populations are denoted by wild type (WT) and escape-variant or mutant (MT).

## 2 Models and Results

As mentioned in the Introduction, there are (at least) three factors that can affect the fitness of an escape-variant: (i) The fitness cost of the mutation; (ii) the extent of selection for the escape-variant in hosts carrying the *relevant* MHC-I allele(s) (i.e. one that presents the wild-type form of CD8 T cell epitope); (iii) the frequency of hosts with the relevant MHC-I allele(s) in the population. We begin with a relatively simple population genetics model that allows us to examine how the interplay between these factors affects the rate of invasion of an escape-variant. This model assumes the selective advantage for the escape-variant in a host with a given set of MHC-I alleles is fixed. We then consider why this assumption may not hold and examine the consequences of relaxing this assumption, using an epidemiological model for the spread of wild type and escape-variant.

### 2.1 Population genetics model

Let us consider the wild type (WT) and one escape-variant (MT). Let *h* represent the set of MHC-I allele(s) that present the focal epitope of WT but not MT, and let *f* equal the frequency of *h*. The MHC-I alleles that present epitopes other than the focal one, *H*, are at frequency (1 − *f*). With the usual assumptions for Hardy-Weinberg equilibrium, we have

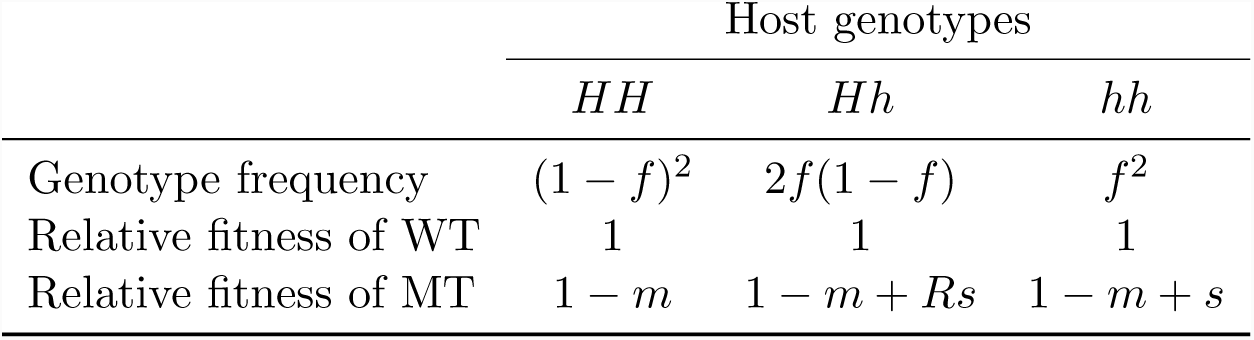

where *m, s, R* denote the fitness cost of MT in all hosts, the selective advantage of MT in hosts of *hh* genotype, and the dominance coefficient of the *h* allele, respectively. Here we have assumed that the WT is equally fit in all genotypes, i.e., while *h* alleles present the focal epitope, the *H* alleles present other epitopes that the virus also carries, so that all alleles confer about the same level of resistance. This is not true for the MT: while the *H* alleles still successfully target its other epitopes, *h* does not recognize its focal epitope. The frequency of the MT in generation *t, q*_*t*_, is given by

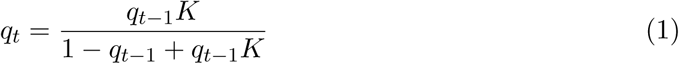

where *K* = (1 − *f*)^2^(1 − *m*) + 2*f* (1 − *f*)(1 − *m* + *Rs*) + *f* ^2^(1 − *m* + *s*) is the mean fitness of MT in the host population.

Equation 1 allows us to examine how the rate of invasion of the MT depends on: *m*, the fitness cost; *f*, the frequency of *h*; *s*, the selective advantage MT accrues in the hosts carrying *h*; and *R*, the heterozygous effect. We first set *R* = 0 on the biological ground that MHC-associated resistance to infections (and susceptibility to autoimmunity) is dominant (16, 17). In Appendix we consider the effect of relaxing this assumption.

In Figure 1, we plot how the rate of invasion depends on the the selective advantage *s* of the MT (*x*-axis) and the frequency *f* of the MHC-I alleles in which MT has a selective advantage (*y*-axis). The rate of invasion is plotted on a log scale – it shows the log10 of the number of generations required for the MT to go from a prevalence of 0.01% to 50% in the host population. Given that the serial interval for influenza is about 3 to 4 days (18), 100-generation corresponds to about one year. In Panel A, we set the fitness cost to 1% (i.e. *m* = 0.01). There is a parameter regime (white region with low *s* and *f*) where the fitness cost is sufficient to prevent MT from invasion. When MT can invade, we see faster invasion when *s* or *f* increase.

**Figure 1:**
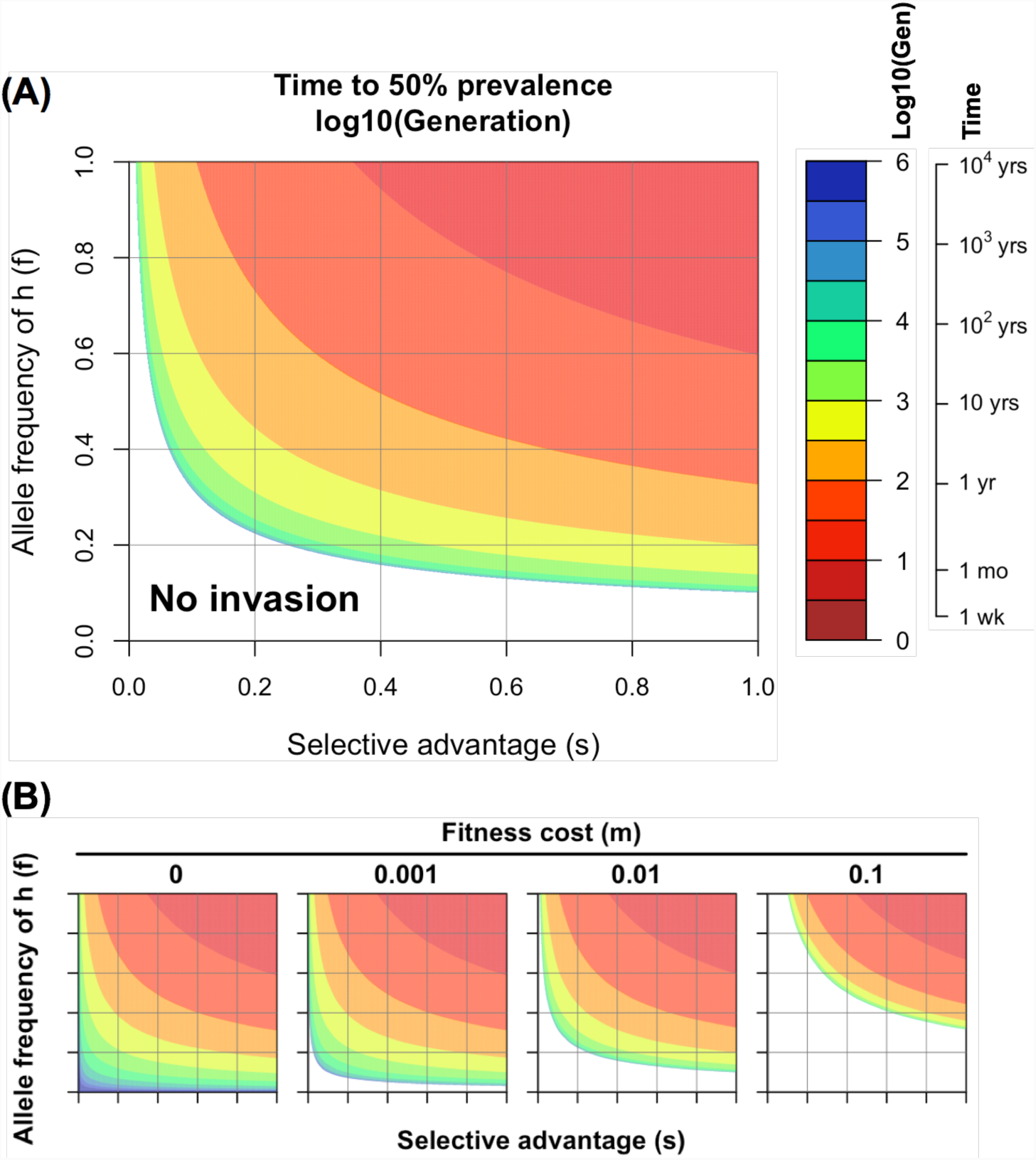
Rate of invasion of a CD8 T cell escape-variant. (A) Contour plots show the number of generations on a log scale required for the escape-variant to increase from 0.01 to 50% prevalence predicted by Equation 1. The approximate time for this to occur is calculated by assuming a serial interval of 3-4 days between infections (i.e. that there are about 100 generations per year). Fitness cost is set to 1% (i.e., *m* = 0.01). (B) We show the time for invasion under different fitness costs, which goes from 0 to 10%. We see that even when the mutant does not have a fitness cost (*m* = 0), it will invade relatively slowly if *s* and *f* are small due to the nature of T cell protection and extent of MHC polymorphism. Ticks on the axes in (B) indicate the same numbers as in (A).

#### Parameter *m*

We show the effect of changing the fitness cost of MT (*m*) from 0 to 10% in Figure 1B. We see that even if the escape-variant does not have any fitness cost (i.e., *m* = 0), it invades relatively slowly when *s* and *f* are small (the region to the bottom left of the leftmost panel of Figure 1B). See Appendix for more details.

#### Parameter *s*

Building on the earlier ideas proposed by Halloran et al. (19), immunity can provide protection by reducing susceptibility of immune individuals to infection (IE_*S*_), as well as by reducing pathology (IE_*P*_) and transmission (IE_*I*_) in infected individuals. While the role of CD8 T cells in providing protection against influenza remains to be fully understood, a number of studies suggest that they play a significant role in reducing pathology (high IE_*P*_). In humans, higher CD8 T cell responses prior to heterosubtypic virus infection are associated with faster viral clearance (6) and fewer symptoms (7). In support of human studies, mouse experiments have shown the cellular immunity induced by H1N1 and/or H3N2 is able to protect the hosts from lethal infection with avian H5N1 or H7N9 (20, 21, 22). CD8 T cells are likely to be less effective in preventing infection (very low IE_*S*_), although they may reduce the viral load during infection (modest IE_*I*_). Consequently, the selection pressure on a virus imposed by *all* CD8 T cell responses, 1 − (1 − IE_*S*_)(1 − IE_*I*_), will be relatively low. Furthermore, since a virus has multiple CD8 T cell epitopes, the selective advantage of a variant that escapes CD8 T cell responses to a single epitope would be considerably smaller (See Appendix for details).

In conclusion, although CD8 T cells may provide some protection against severe pathology, escape-variants having a mutated CD8 T cell epitope are unlikely to have much selective advantage, even in the hosts with the relevant MHC-I that presents the wild-type epitope. In other words, we expect *s* to be small.

#### Parameter *f*

Escape-variants will only accrue an advantage in individuals of *hh* genotype. For example, the R384G escaping mutation on the NP_383-391_ epitope is advantageous in the hosts who carry B*08:01 and/or B*27:05 but not in those who carry other alleles. In Figure 2, we show the distribution of experimentally-verified CD8 T cell epitopes derived from the nucleoprotein of human IAV retrieved from the Immune Epitope Database. It is clear that no single epitope is presented by all human leucocyte antigen (HLA, the human version of MHC) alleles, nor is there a single HLA allele presenting all epitopes. With this information and the frequencies of HLA alleles based on the National Marrow Donor Program (NMDP) dataset, we estimated the fraction of host population where the mutation at each amino acid residue would confer a selective advantage (top panel of Figure 2) (see Appendix for details). Typically, mutations confer selective advantage to the virus in only a small fraction (less than 10%) of host population.

**Figure 2:**
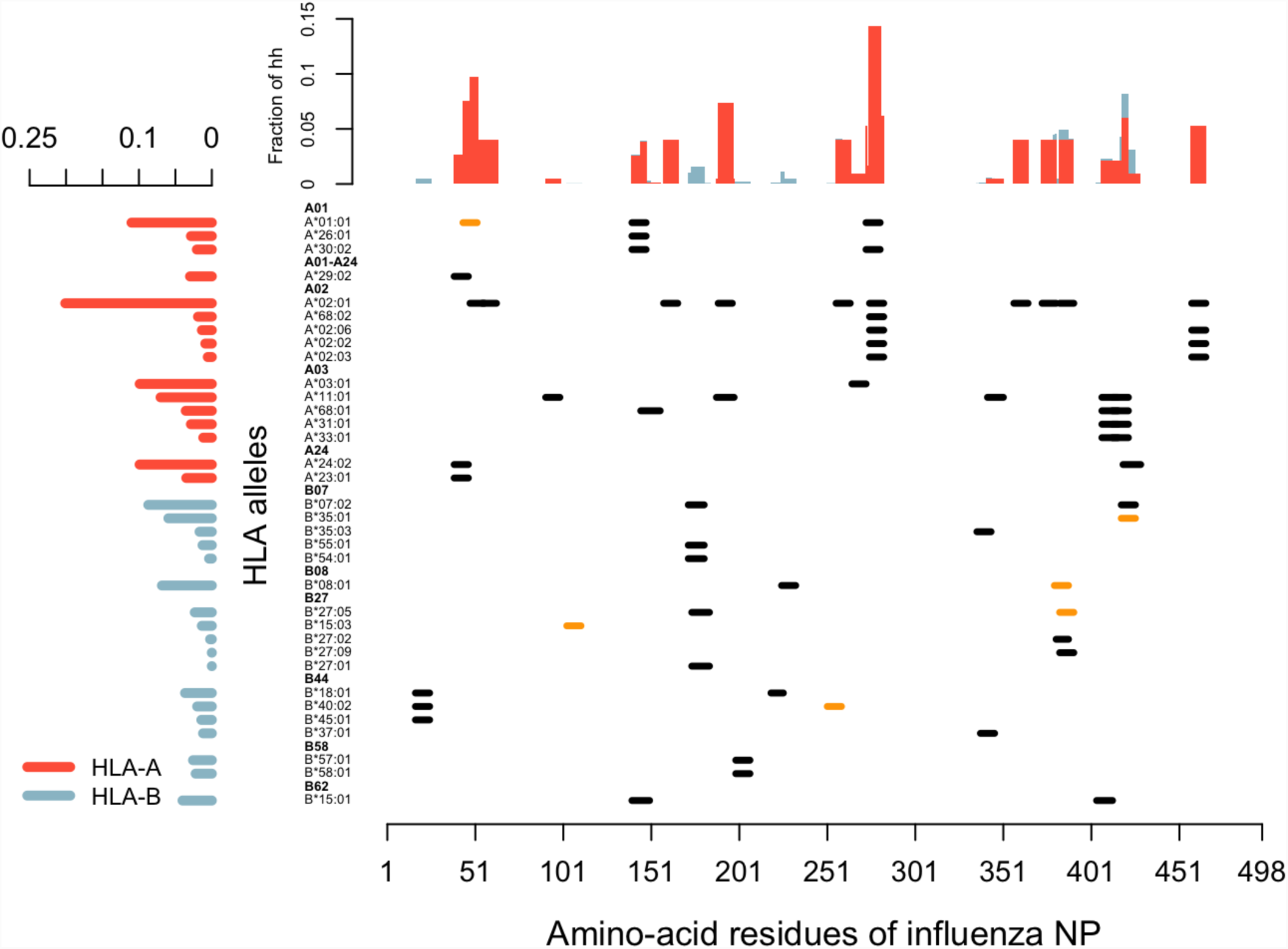
Distribution of CD8 T cell epitopes derived from the nucleoprotein of human IAV. In the main panel, each segment represents one unique CD8 T cell epitope derived from nucleoprotein of human IAV, aligned with its relevant HLA allele. Epitopes that have empirically verified escape-mutants reported are labeled in orange. The bar graph in left panel shows the weighted average frequency of the alleles, which are grouped into 11 supertypes (highlighted in boldface). The bar graph in the top panel shows the fraction of the population in which the virus with a mutation in the corresponding amino acid has a selective advantage.

#### Integrating parameters *s, f*, and *m*

As mentioned in the Introduction, epistatic interactions allow the virus to potentially generate escape-variants to a given CD8 T cell epitope without engendering a substantial fitness cost. Our results show that, since *s* and *f* are small, these escape-variants will spread very slowly in the host population. For example, when *f* = 0.1 (i.e. the escape-variant has an advantage in about 1% of the infections, who are of *hh* genotype (*f* ^2^ = 0.01)), an escape-variant would spend 50 to 100 years reaching 50% prevalence, even if its fitness is 20% more than the wild type (i.e. *s* = 0.2) in the hosts with relevant HLA and no fitness cost is accompanied (i.e., *m* = 0).

In conclusion, even if mutations that allow the virus to escape CD8 T cells specific for a given epitope have little or no fitness cost, escape-variants will only increase in frequency very slowly.

### 2.2 Epidemiological models

#### Compensatory immunity reduces the selective advantage of mutants over time

The population genetics framework described above assumes that the fitness of an escape-variant virus depends on host genotype and does not change over time. In particular, we assume that the fitness of escape-variant (MT) relative to the wild-type (WT) equals (1 − *m* + *s*) in the hosts of the *hh* genotype and (1 − *m*) in the hosts of other genotypes (*HH* and *Hh*). However, in a host of *hh* genotype who has been infected by MT, recovered, and moved to the susceptible category (due to antigenic drift), we might expect the selective advantage (*s*) of MT to decrease. This decline in *s* could arise for at least two biological reasons. First, the mutated epitope is still presented by the MHC-I; the mutation simply changes the configuration of the epitope recognized by the CD8 T-cell receptor (12). In this case, the mutant epitope may induce a new set of CD8 T cells. Second, the lack of CD8 T cell response to one epitope could result in compensatory increases in responses to other epitopes.

We show the fitness of WT or MT infections in the hosts of different genotypes in Figure 3. In hosts of *HH* and *Hh*, the fitnesses of WT and MT are 1 and (1 − *m*), the same as in population genetics model. The fitness of MT in hosts of *hh* that are infected with MT for the first time is (1 − *m* + *s*) as described earlier. After these hosts recover and regain susceptibility due to antigenic drift, the fitness of MT following reinfection with MT becomes (1 − *m* + *s*(1 − *c*)), where *c* denotes the extent of compensatory CD8 T cell responses. Parameter *c* ranges from 0 to 1, with 0 corresponding to no compensatory increase in responses to other epitopes and 1 corresponding to full compensation. The range for the fitness of MT in hosts of *hh* with prior infections with MT is shown by the shaded region in Figure 3A.

**Figure 3:**
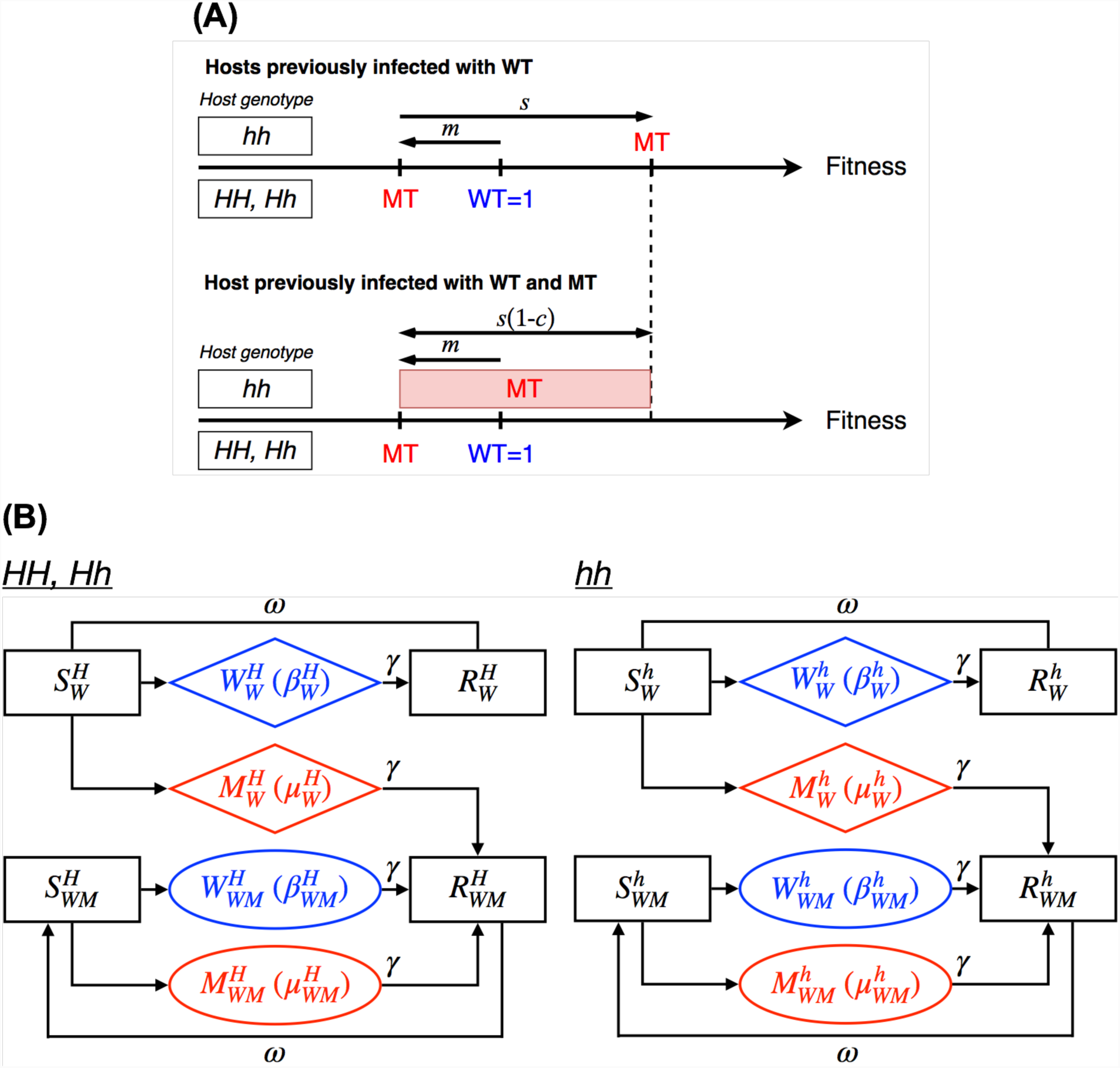
(A) Compensatory immunity alters the fitness of escape-variant infection in the *hh* individuals. We show the viral fitness in the hosts of different genotypes and infection histories, when they are infected with the wild type (WT) or escape-mutant (MT). The WT is shown in blue and MT in red. The fitness of WT infections is 1 in all hosts. The fitness of MT in *HH* and *Hh* hosts is (1 − *m*) while in *hh* hosts who are infected for the first time is (1 − *m* + *s*). Subsequent MT infections of *hh* hosts result in lower viral fitness (1 − *m* + *s*(1 − *c*)) due to compensatory immunity, which reduces the selective advantage by *c*. (B) Diagram illustrating the epidemiology of infections with WT (*W* shown in blue) and MT (*M* shown in red). Susceptible (*S*) and immune (*R*) hosts are indicated by *S* and *R*, respectively. The host genotype is indicated by the superscript *j* (*h* for *hh* and *H* for *HH/Hh*), and the prior infection status is indicated by the subscript (*W* for WT and *M* for MT).

### Epidemiology of infections with wild-type and escape-variant viruses

We use a simple epidemiological model to describe the changes in frequencies of susceptible (*S*), infected (*W* and *M* for WT and MT infections) and immune (*R*) hosts. The subscript to *S, W, M*, and *R* populations indicates the viruses these hosts have been exposed to in the past. Individuals can be infected multiple times during their lifetime due to antigenic drift at antibody epitopes (23). We incorporate this by letting individuals move from the immune (*R*) to susceptible (*S*) compartments at rate *ω* (24).

We consider the epidemiology of WT infections in individuals of different genotypes, where the right section of Figure 3B is hosts of *hh* genotype and the left section is hosts of *HH* and *Hh* genotypes. Prior to the introduction of MT, we assume that the WT is circulating and hosts have CD8 T cell immunity to the wild-type epitope. Due to antigenic drift, individuals typically get reinfected with a drifted strain every 5-10 years (23), and we choose the rate of loss of immunity corresponding to this duration (*ω* = 5 × 10^-4^/day ≈ 5.5/year). We begin the simulations with WT infections at equilibrium.

Now, on the introduction of MT, the MT has fitness (1 − *m*) or (1 − *m* + *s*) in the MT-infected hosts of the *HH*/*Hh* or *hh* genotypes, respectively. MT-infected individuals move to the immune category with a subscript of *WM* (e.g. *R*_*WM*_). Individuals in *R*_*WM*_ become susceptible due to antigenic drift in the virus and move to *S*_*WM*_. When individuals in *S*_*WM*_ are infected with the WT, they move to *W*_*W* *M*_ and the WT has fitness 1, while when individuals in *S*_*WM*_ are infected with the MT, they move to the *M*_*W* *M*_ and the MT has fitness (1*-m*+*s*(1*-c*)). For simplicity, we incorporate the fitness in the transmissibility parameter (*β*). Equations are shown in the Appendix.

In Figure 4, we explore how the escape-variant (MT) spreads through the host population following its introduction. In particular, we focus on how compensatory immunity changes the outcomes predicted by the population genetics models. In Panel 4A, we chose a simple scenario where the MT has a very small fitness cost (*m* = 0.001), a 5% selective advantage (*s* = 0.05) in 10% of the population (*f* ^2^ = 0.1), and compensatory immunity reduces the selective advantage by 90% (*c* = 0.9). In this scenario, we see that the MT now only transiently invades, and compensation in host immunity causes the frequency of MT to decline as the population-level immunity against the MT increases. In Panel 4B, we explore the consequences of changing the extent of compensation (*c*). We see that the initial rate of invasion is very similar to what is predicted by the population genetics model. However, once the MT has spread through the population, the outcome depends strongly on the extent of compensatory immunity described by the parameter *c*. If *c* is small, then the MT goes to fixation in a manner similar to that of the population genetics model. If *c* exceeds a threshold value *c\** given by

**Figure 4:**
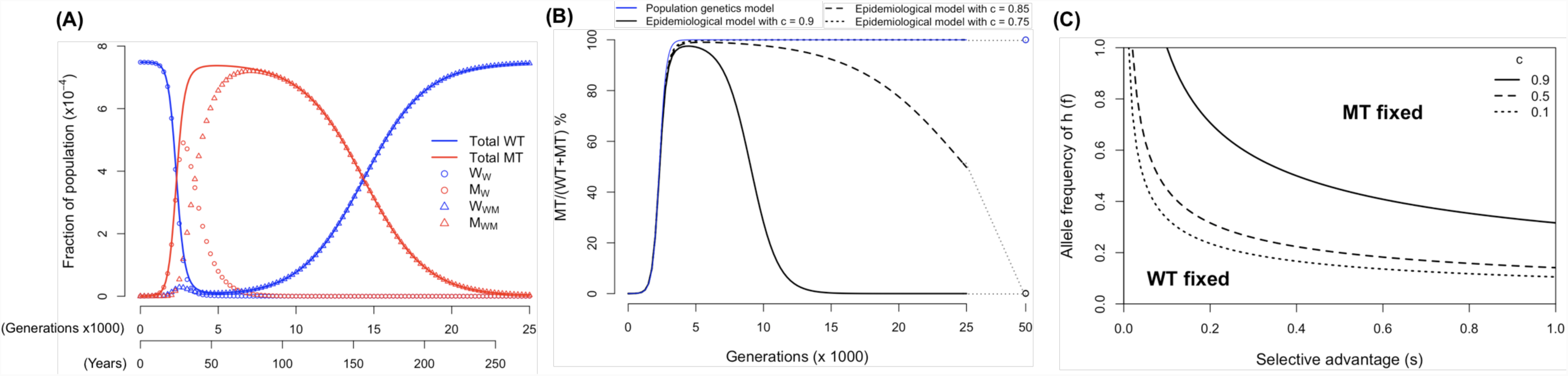
(A) The frequency of wild-type (WT) and escape-variant (MT) infections as a function of time. When CD8 T cell responses are compensated and thus decreases the selective advantage, the MT only invades transiently and goes extinct in the long run. (*m* = 0.001, *s* = 0.05, *f* ^2^ = 0.1, *c* = 0.9). (B) The MT prevalence predicted by epidemiological model with different degrees of compensation (black lines), compared to the population genetics model (blue line). In all scenarios, the initial invasions of the MT (before it reaches 50% prevalence) are similar. After it reaches 50%, if the degree of compensation is smaller than the threshold (in our parameter setting, *c\** = 0.8. See Equation 2), the MT goes to fixation only slightly slower than what predicted by the population genetics model. In contrast, if the degree of compensation is higher than the threshold, the MT invades transiently and becomes extinct in the long run. (C) The parameter regimes of *s* and *f* where either MT or WT becomes fixed (and the other becomes extinct). We see that as the degree of compensation rises, the parameter regime where MT becomes fixed shrinks.

**Figure 5:**
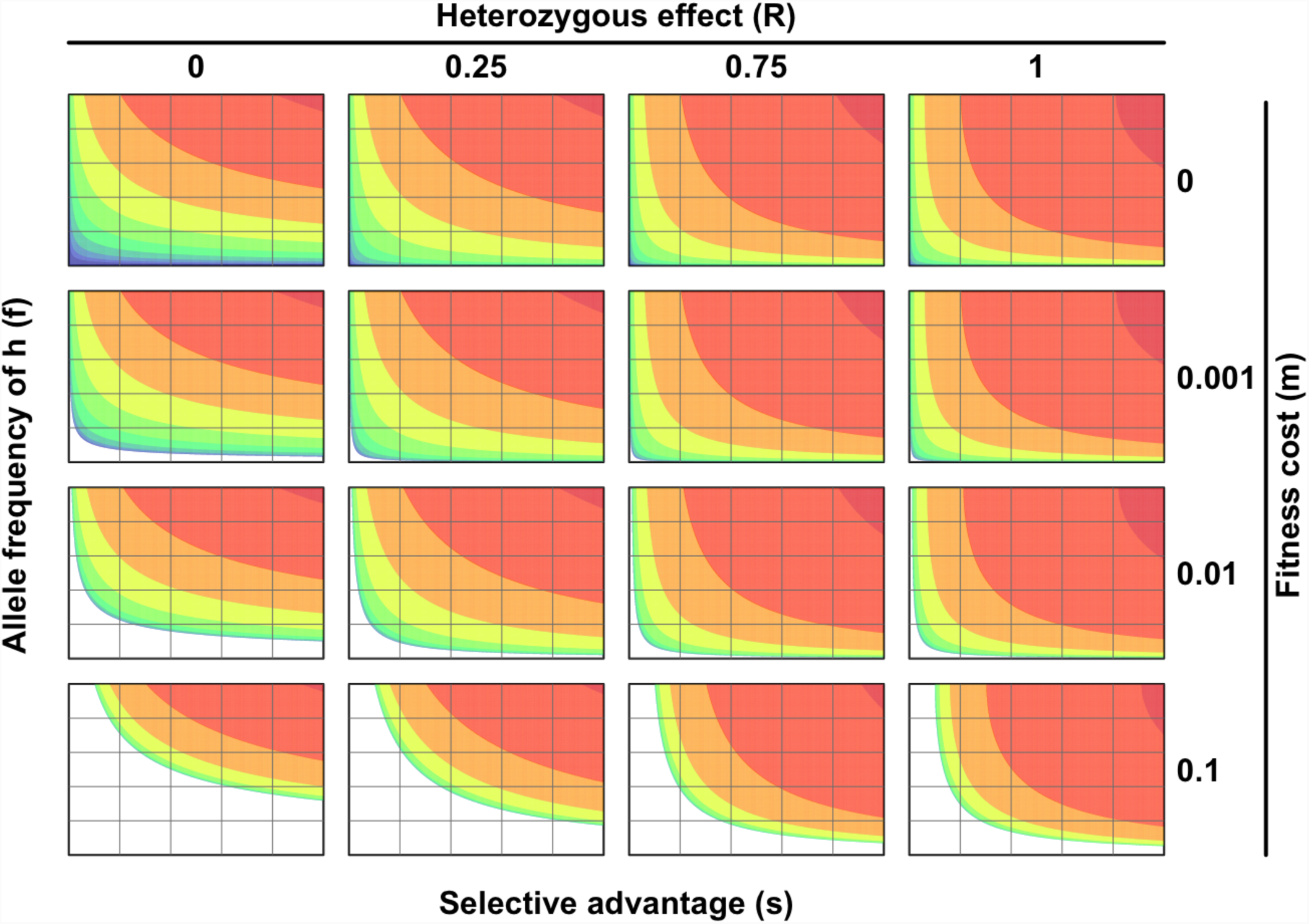
Rate of invasion of an escape-variant when the assumption of *R* = 0 is relaxed.

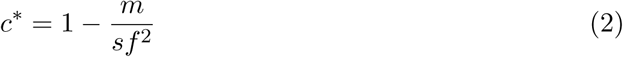

then the MT only transiently invades but declines to extinction in the long run. The competitive exclusion between WT and MT infections is shown in Panel 4C, where we plot how the outcome depends on *s, f*, and *c*. Incorporating fitness parameters into the duration of infection gives similar results (results not shown).

In summary, the results of the epidemiological models show that the initial rate of invasion of the escape-variant is similar to that in the population genetics model described by Equation 1. However, at later time points, compensatory immunity reduces the rate of invasion. If the extent of CD8 T cell immunity to the escape-variant is sufficiently high, the outcome may even reversed if the overall selective advantage does not surmount fitness cost.

## 3 Discussion

We consider the question why CD8 T cell epitopes of human IAV are conserved. The answer to this question is relevant to the development of ‘universal’ CD8 T cell-inducing vaccines against influenza, and whether the virus is likely to evolve to escape the vaccine-induced immunity. Despite the wealth of empirical data showing conservation of CD8 T cell epitopes, the evolutionary mechanisms responsible for this conservation are not well known. One possibility that has been widely considered is that the nucleoprotein and matrix-1 protein, which harbor the bulk of CD8 T cell epitopes, are under strong constraints (13, 14). In this view, mutations in CD8 T cell epitopes would have a high fitness cost, or be relatively inaccessible due to epistatic changes needed for escape-variants to have high fitness. In this study, we proposed two other mechanisms that could impact the fitness of viruses that escape CD8 T cell recognition. The first one is that escape from CD8 T cell responses against a single epitope provides only a relatively small selective advantage to the escape-variant. The second one is that polymorphism in the MHC-I genes restricts this small advantage to only a small fraction (typically less than 10%) of individuals in the population – individuals with other alleles would not present this epitope but present other epitopes. We show that even if there is a minimal fitness cost to having a mutation in a CD8 T cell epitope, the latter two factors will result in a very low rate of invasion of the CD8 T cell escape-variant.

The conclusion of this study may seem to contradict the rapid invasion of the mutation at the 384-th amino-acid residue of the nucleoprotein. This mutation alters the NP_383-391_ epitope presented by HLA-B*27:05 and NP_380-388_ epitope presented by HLA-B*08:01. The wild-type sequence has an arginine (R), which forms an anchor residue at this site, and all 16 viruses isolated and sequenced during the 1992-1993 epidemic season had the wild-type sequence (25). An arginine-to-glycine mutation at this site (R384G) abrogates the MHC-I binding and prevents antigen presentation, and this mutation rapidly swept through the population in the 1993-4 epidemic season, where all 56 virus isolates had G at this residue (25). Gog et al. (26) suggested that the rapid fixation of the R384G mutation was due to a combination of a longer duration of infection that slows its decline compared to the wild-type over the summer and stochastic events. We propose an alternative hypothesis – a selective advantage of the R384G mutation is not required and hitchhiking of a randomly generated mutant would be sufficient to explain the data. The rapid invasion of R384G temporally matches the transition from BE92 to WU95 antigenic clusters (27), suggesting that it could have hitchhiked with the antigenically drifted WU95 strain.

A recent analysis of the number of CD8 T cell epitopes in circulating strains of influenza showed that they gradually decline over a timescale of decades (28). For example, the H3N2 subtype had 84 experimentally-confirmed epitopes per virus in 1968, and this number declined to 64 in 2015. A decline in the number of epitopes could be partially due to the bias in identification of epitopes, as new epitopes may not be identified and included. The observed decline rather than drift in the number of confirmed epitopes may arise because of a slight selective advantage of the escape-variants over the wild-type. This study suggests that the gradual escape of the virus from a CD8 T cell vaccine may be possible.

We have intentionally used simple models – this is because the empirical data does not include accurate measurements of many of the key parameters that govern the generation and spread of virus escape-variants. In these circumstances the results of simpler models are typically more robust than those of complex models (29). This study identifies the importance of measuring parameters such as the fitness of the wild-type and escape-variants in hosts, who have been previously infected with wild-type and both wild-type and escape-variants. Aspects that might be included in more refined models include: The waning of CD8 T cell immunity over time, particularly due to the loss of resident memory cells from the respiratory tract (30, 31), and the effect of stochasticity.

There are several differences in the ability of the influenza virus to evolve in response to antibody and CD8 T cell immunity. First, antibody immunity can generate sterilizing immunity (prevent infection with a matched virus strain) and thus generates substantial selection for antibody-escape-variants. In contrast, CD8 T cell immunity to influenza does not prevent infection and thus generates less selective advantage to a CD8 T cell-escape-variant. Second, an antibody-escape-variant will gain a selective advantage in the majority of individuals with antibodies to the wild-type strain while a CD8 T cell-escape-variant will have a selective advantage only in individuals of a particular MHC-I genotype. Both these factors contribute to antibody rather than CD8 T cell immunity driving antigenic drift in influenza.

Vaccination strategies that boost the CD8 T cell response may contribute to the development of broadly protective influenza vaccines. In this paper, we focus on whether these vaccine strategies will rapidly select for virus escape-variants at CD8 T cell epitopes, compromising the effectiveness of the vaccine. We show that this is unlikely to be the case. Although it is generally viewed as a potential limitation that these vaccines may not completely prevent infection, this fact, together with MHC polymorphism, greatly reduces the selection pressure on the virus. Consequently, it may take a much longer duration for the virus to evolve and escape vaccine-induced CD8 T cell immunity.

## Appendix

### Population genetics model

Equation 1 can be rearranged into

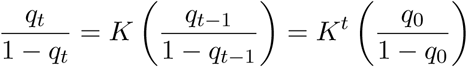

We then express *t*, the number of generation required for the MT to reach *q*_*t*_, by

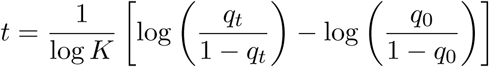

The escape-variant can invade when *K* > 1, i.e.,

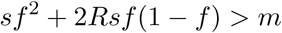

### Quantifying selection pressure on virus

We define three types of immunity effectiveness (IE) as follows:

- IE_*S*_ is the probability that an individual who have influenza-specific CD8 T cells does not get infected upon a contact with an infectious individual
- IE_*P*_ is the probability that an infected who have influenza-specific CD8 T cells does not develop symptoms.
- IE_*I*_ as the probability that an infected who have influenza-specific CD8 T cells does not spread the virus.

The selection pressure on virus (*S*) is formulated by

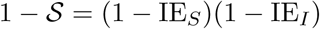

The viral fitness after selection, 1 − *S*, is the probability of transmission within the host population who have influenza-specific CD8 T cells, and can be expressed as the probability that an immune individual gets infected (1 − IE_*S*_) and spreads the virus (1 − IE_*I*_).

### Map of CD8 T cell epitope on influenza nucleoprotein

Epitope dataset was retrieved from Immune Epitope Database (www.iedb.org). We searched for MHC class I-restricted linear epitope of influenza A virus (ID: 11320, FLUAV) in humans with at least one positive T cell assay. We retrieved 1,220 records from IEDB, of which 514 were derived from NP. After excluding the records longer than 12 amino-acid residues or with no HLA allele information available, records with the same amino-acid sequence, with different sequences but at the same location of NP and presented by the same HLA allele, or nested under a longer record, were combined into one ‘unique’ epitope. In total, 64 unique epitopes were identified. Escaping mutations were identified from the literatures (9, 12).

HLA allele dataset reported by National Marrow Donor Program (NMDP) was retrieved from The Allele Frequency Net Database (www.allelefrequencies.net). We included all the alleles that have been reported to present at least one epitope in the epitope dataset, and calculated the average frequency weighted by sample sizes. In addition, since the alleles in one HLA supertype prefer amino acid with similar chemical property at certain residues of the epitopes, we grouped the HLA alleles based on the classification proposed by Sette et al. (32).

With the epitope and HLA allele datasets, we estimate the fraction of host population affected on each amino-acid residue given that there is an escaping mutation, assuming (1) the escaping effect is recessive and (2) the alleles of one locus are under HWE. For a particular residue included in a number of epitopes, we denote the collection of unique alleles that can present these epitopes, {*h*_1_, *h*_2_, *· · ·, h*_*m*_}, by *h*. The mutant is able to escape only from the hosts carrying both alleles from *h*, and the estimated probability of escape is given by

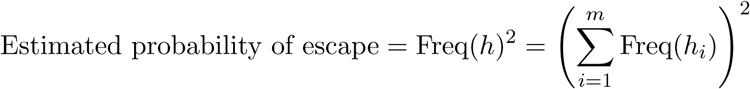

For example, the T147 residue is included in three epitopes: NP140-148 (bound by A*01:01, A*26:01, and A*30:02), NP140-150 (bound by B*15:01), and NP145-156 (bound by A*68:01). Suppose an escaping mutation on T147 results in escape from all of the alleles, the estimated probability of this mutant escaping from a host is

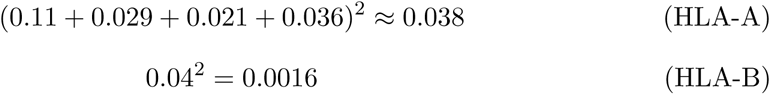

Although the Assumption (1) has not been tested in the context of influenza infection, studies on the genetic factors of autoimmune diseases may provide indirect support. Risk alleles associated with multiple sclerosis and type 1 diabetes mellitus have dominant genetic predisposition to the diseases (16, 17). It implies that one allele is enough to present antigen and activate autoreactive CD4 T cell; conversely, it implies that it is not enough for a virus to escape from detection if only one allele is escaped.

### ODE system and the equilibrium

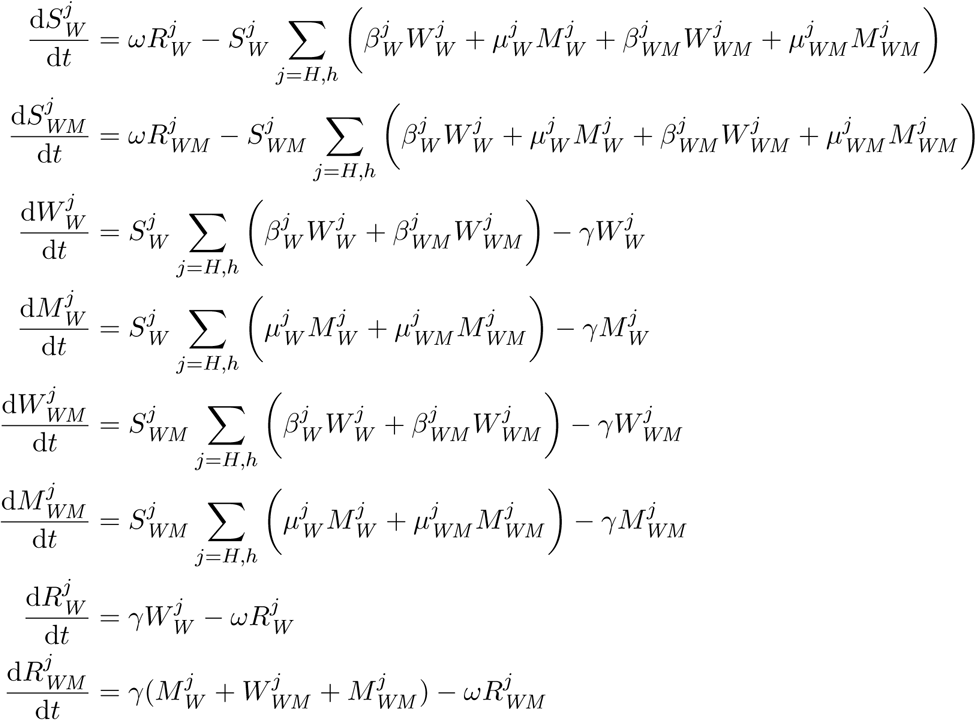

where *j* = *h* denotes the genotype of *hh* and *j* = *H* denotes the genotypes of *HH* and *Hh*. We started simulations from the equilibrium of WT infection, i.e.,

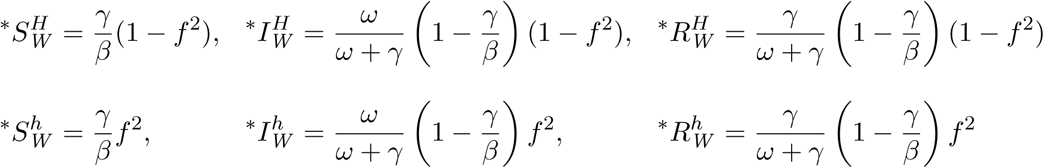

where 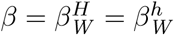. Values of parameters are listed in Table 1.

**Table 1:** Model parameters

| Parameter | Symbol | Value |
| --- | --- | --- |
| Transmission rate of $W_W^H$ ($\text{day}^{-1}$)<sup>†</sup> | $\beta_W^H$ | 0.4 |
| Recovery rate ($\text{day}^{-1}$) | $\gamma$ | 0.25 |
| Drifting rate ($\text{day}^{-1}$) | $\omega$ | $5 \times 10^{-4}$ |

<sup>†</sup>See Table 2 for the setup of other transmission parameters.

**Table 2:** Transmissibility according to genotypes and immune status Compartment Symbol Value

| Compartment | Symbol | Value |
| --- | --- | --- |
| $M_W^H$ | $\mu_W^H$ | $\beta_W^H(1 - m)$ |
| $W_W^h$ | $\beta_W^h$ | $\beta_W^H$ |
| $M_W^h$ | $\mu_W^h$ | $\beta_W^H(1 - m + s)$ |
| $W_{WM}^H$ | $\beta_{WM}^H$ | $\beta_W^H$ |
| $M_{WM}^H$ | $\mu_{WM}^H$ | $\beta_W^H(1 - m)$ |
| $W_{WM}^h$ | $\beta_{WM}^h$ | $\beta_W^H$ |
| $M_{WM}^h$ | $\mu_{WM}^h$ | $\beta_W^H(1 - m + (1 - c)s)$ |

## References

1. Iuliano AD, Roguski KM, Chang HH, Muscatello DJ, Palekar R, Tempia S, Cohen C, Gran JM, Schanzer D, Cowling BJ, Wu P, Kyncl J, Ang LW, Park M, Redlberger-Fritz M, Yu H, Espenhain L, Krishnan A, Emukule G, van Asten L, da Silva SP, Aungkulanon S, Buchholz U, Widdowson M-A, Bresee JS, and the Global Seasonal Influenza-associated Mortality Collaborator Network. 2017. Estimates of global seasonal influenza-associated respiratory mortality: A modelling study. Lancet 391(10127):1285–1300.

2. Hensley SE, Das SR, Bailey AL, Schmidt LM, Hickman HD, Jayaraman A, Viswanathan K, Raman R, Sasisekharan R, Bennink JR, and Yewdell JW. 2009. Hemagglutinin receptor binding avidity drives influenza A virus antigenic drift. Science 326(5953):734–736.

3. Soema PC,Kompier R,Amorij JP,and Kersten GF. 2015. Current and next generation influenza vaccines: Formulation and production strategies. Eur J Pharm Biopharm 94:251–263.

4. Krammer F and Palese P. Advances in the development of influenza virus vaccines. 2015. Nat Rev Drug Discov 14(3):167–182.

5. Clemens E, van de Sandt C, Wong S, Wakim L, and Valkenburg S. 2018. Harnessing the power of T cells: The promising hope for a universal influenza vaccine. Vaccines (Basel) 6(2):18.

6. McMichael AJ, Gotch FM, Noble GR, and Beare PA. 1983. Cytotoxic T-cell immunity to influenza. N Engl J Med 309(1):13–17.

7. Sridhar S, Begom S, Bermingham A, Hoschler K, Adamson W, Carman W, Bean T, Barclay W, Deeks JJ, and Lalvani A. 2013. Cellular immune correlates of protection against symptomatic pandemic influenza. Nat Med 19(10):1305–1312.

8. Liang S, Mozdzanowska K, Palladino G, and Gerhard W. 1994. Heterosubtypic immunity to influenza type A virus in mice. Effector mechanisms and their longevity. J Immunol 152:1653–1661.

9. Quinones-Parra S, Grant E, Loh L, Nguyen THO, Campbell K-A, Tong SYC, Miller A, Doherty PC, Vijaykrishna D, Rossjohn J, Gras S, and Kedzierska K. 2014. Preexisting CD8+ T-cell immunity to the H7N9 influenza A virus varies across ethnicities. Proc Natl Acad Sci U S A 111(3):1049–1054.

10. Bhatt S, Holmes EC, and Pybus OG. 2011. The genomic rate of molecular adaptation of the human influenza A virus. Mol Biol Evol 28(9):2443–2451.

11. Machkovech HM, Bedford T, Suchard MA, and Bloom JD. 2015. Positive selection in CD8+ T-cell epitopes of influenza virus nucleoprotein revealed by a comparative analysis of human and swine viral lineages. J Virol 89(22):11275–11283.

12. Rimmelzwaan GF, Kreijtz JHCM, Bodewes R, Fouchier RAM, and Osterhaus ADME. 2009. Influenza virus CTL epitopes, remarkably conserved and remarkably variable. Vaccine 27(45):6363–6365.

13. Berkhoff EGM, de Wit E, Geelhoed-Mieras MM, Boon ACM, Symons J, Fouchier RAM, Osterhaus ADME, and Rimmelzwaan GF. 2005. Functional constraints of influenza A virus epitopes limit escape from cytotoxic T lymphocytes. J Virol 79(17):11239–11246.

14. Berkhoff EGM, de Wit E, Geelhoed-Mieras MM, Boon ACM, Symons J, Fouchier RAM, Osterhaus ADME, and Rimmelzwaan GF. 2006. Fitness costs limit escape from cytotoxic T lymphocytes by influenza A viruses. Vaccine 24(44-46):6594–6596.

15. Gong LI, Suchard MA, and Bloom JD. 14 May 2013, posting date. Stability-mediated epistasis constrains the evolution of an influenza protein. Elife 2:e0063doi:10.7554/eLife.00631.

16. Simmonds M and Gough S. 2007. The HLA region and autoimmune disease: Associations and mechanisms of action. Curr Genomics 8(7):453–465.

17. Cruz-Tapias P, Castiblanco J, and Anaya J-M. 2013. HLA association with autoimmune diseases, pp. 271–28In Anaya JM, Shoenfeld Y, Rojas-Villarraga A, Levy RA, Cervera R (ed.), Autoimmunity: From Bench to Bedside. El Rosario University Press, Bogota (Columbia).

18. Cowling BJ, Fang VJ, Riley S, Peiris JSM, and Leung GM. 2009. Estimation of the serial interval of influenza. Epidemiology 20(3):344–347.

19. Halloran ME, Struchiner CJ, and Longini IM. 1997. Study designs for evaluating different efficacy and effectiveness aspects of vaccines. Am J Epidemiol 146(10):789–803.

20. O’Neill E, Krauss SL, Riberdy JM, Webster RG, and Woodland DL. 2000. Heterologous protection against lethal A/HongKong/156/97 (H5N1) influenza virus infection in C57BL/6 mice. J Gen Virol 81(Pt 11):2689–2696.

21. Kreijtz JHCM, Bodewes R, van den Brand JMA, de Mutsert G, Baas C, van Amerongen G, Fouchier RAM, Osterhaus ADME, and Rimmelzwaan GF. 2005. Infection of mice with a human influenza A/H3N2 virus induces protective immunity against lethal infection with influenza A/H5N1 virus. Vaccine 27(36):4983–4989.

22. McMaster SR, Gabbard JD, Koutsonanos DG, Compans RW, Tripp RA, Tompkins SM, and Kohlmeier JE. 11 Feb 2015, posting date. Memory T cells generated by prior exposure to influenza cross react with the novel H7N9 influenza virus and confer protective heterosubtypic immunity. PLoS One 10(2):e011572doi:10.1371/journal.pone.0115725.

23. Kucharski AJ, Lessler J, Read JM, Zhu H, Jiang CQ, Guan Y, Cummings DAT, and Riley S. 2013 Mar 2015, pposting date. Estimating the life course of influenza A(H3N2) antibody responses from cross-sectional data. PLoS Biol 13(3):e100208doi:10.1371/journal.pbio.1002082.

24. Pease CM. An evolutionary epidemiological mechanism, with applications to type A influenza. 1987. Theor Popul Biol, 31(3):422–452.

25. Voeten JTM, Bestebroer TM, Nieuwkoop NJ, Fouchier RAM, Osterhaus ADME, and Rimmelzwaan GF. 2000. Antigenic drift in the influenza A virus (H3N2) nucleoprotein and escape from recognition by cytotoxic T lymphocytes. J Virol 74(15):6800–6807.

26. Gog JR, Rimmelzwaan GF, Osterhaus ADME, and Grenfell BT. Population dynamics of rapid fixation in cytotoxic T lymphocyte escape mutants of influenza A. 2003. Proc Natl Acad Sci U S A 100(19):11143–11147.

27. Smith DJ, Lapedes AS, de Jong JC, Bestebroer TM, Rimmelzwaan GF, Osterhaus ADME, and Fouchier RAM. 2004. Mapping the antigenic and genetic evolution of influenza virus. Science 305(5682):371–376.

28. Woolthuis RG, van Dorp CH, Kesmir C, de Boer RJ, and van Boven M. 2016. Longterm adaptation of the influenza A virus by escaping cytotoxic T-cell recognition. Sci Rep 6:33334.

29. Hilborn R and Mangel M. 1997. The ecological detective: Confronting models with data. Princeton University Press, Princeton.

30. Wu T, Hu T, Lee Y-T, Bouchard KR, Benechet A, Khanna K, and Cauley LS. 2014. Lung-resident memory CD8 T cells (TRM) are indispensable for optimal cross-protection against pulmonary virus infection. J Leukoc Biol 95(2):215–224.

31. Zarnitsyna VI, Handel A, McMaster SR, Hayward SL, Kohlmeier JE, and Antia R. 9 May 2016, posting date. Mathematical model reveals the role of memory CD8 T cell populations in recall responses to influenza. Front Immunol, 7:165. doi:10.3389/fimmu.2016.00165.

32. Sidney J, Peters B, Frahm N, Brander C, and Sette A. 2008. HLA class I supertypes: A revised and updated classification. BMC Immunol 9:1.

